# An Engineered Nanocomposite Copper Coating with Enhanced Antibacterial Efficacy

**DOI:** 10.1101/2022.04.28.489879

**Authors:** Davood Nakhaie, Teresa C. Williams, Billie Velapatino, Elizabeth A. Bryce, Marthe K. Charles, Edouard Asselin, Amanda M. Clifford

**Affiliations:** Department of Materials Engineering, The University of British Columbia, Vancouver, Canada; Division of Medical Microbiology, Department of Pathology and Laboratory Medicine, Vancouver Coastal Health, Vancouver, Canada; Department of Pathology and Laboratory Medicine, The University of British Columbia, Vancouver, Canada

**Author notes:** Corresponding Author: Amanda M. Clifford, Department of Materials Engineering, The University of British Columbia, 6350 Stores Rd, Vancouver, BC V6T 1Z4, Canada.

**Keywords:** Electrophoretic Deposition, Copper, Antibacterial Surfaces, Nanostructured Coatings

## Abstract

Contaminated surfaces are a major source of nosocomial infection. To reduce microbial bioburden and surface-based transmission of infectious disease, the use of antibacterial and self-sanitizing surfaces, such as copper (Cu), is being explored in clinical settings. Cu has long been known to have antimicrobial activity. However, Gram-positive microorganisms, a class that includes pathogens commonly responsible for hospital-acquired infection such as *Staphylococcus aureus* and *Clostridioides difficile*, are more resilient to its biocidal effect. Inspired by inherently bactericidal nanostructured surfaces found in nature, we have developed an improved Cu coating, engineered to contain nanoscale surface features and thus increase its antibacterial activity against a broader range of organisms. In addition, we have established a new method for facilitating the rapid and continuous release of biocidal metal ions from the coating, through incorporation of an antibacterial metal salt (ZnCl_2_) with a lower reduction potential than Cu. Electrophoretic deposition (EPD) was used to fabricate our coatings, which serves as a low-cost and scalable route for modifying existing conductive surfaces with complex shape. By tuning both the surface morphology and chemistry, we were able to create a nanocomposite Cu coating that decreased the microbial bioburden of Gram-positive *S.aureus* by 94% compared to unmodified Cu.

**Table of Contents:** Antimicrobial copper (Cu) products are being deployed in clinical settings to decrease microbial bioburden and prevent surface-based transmission of infectious disease. However, Gram-positive bacteria demonstrate increased resistance to Cu’s biocidal effects. To improve Cu’s antibacterial efficacy against Gram-positive bacteria, we have developed a hydrophobic Cu coating with cytotoxic nanotopography that facilitates the rapid and continuous release of biocidal metal ions.

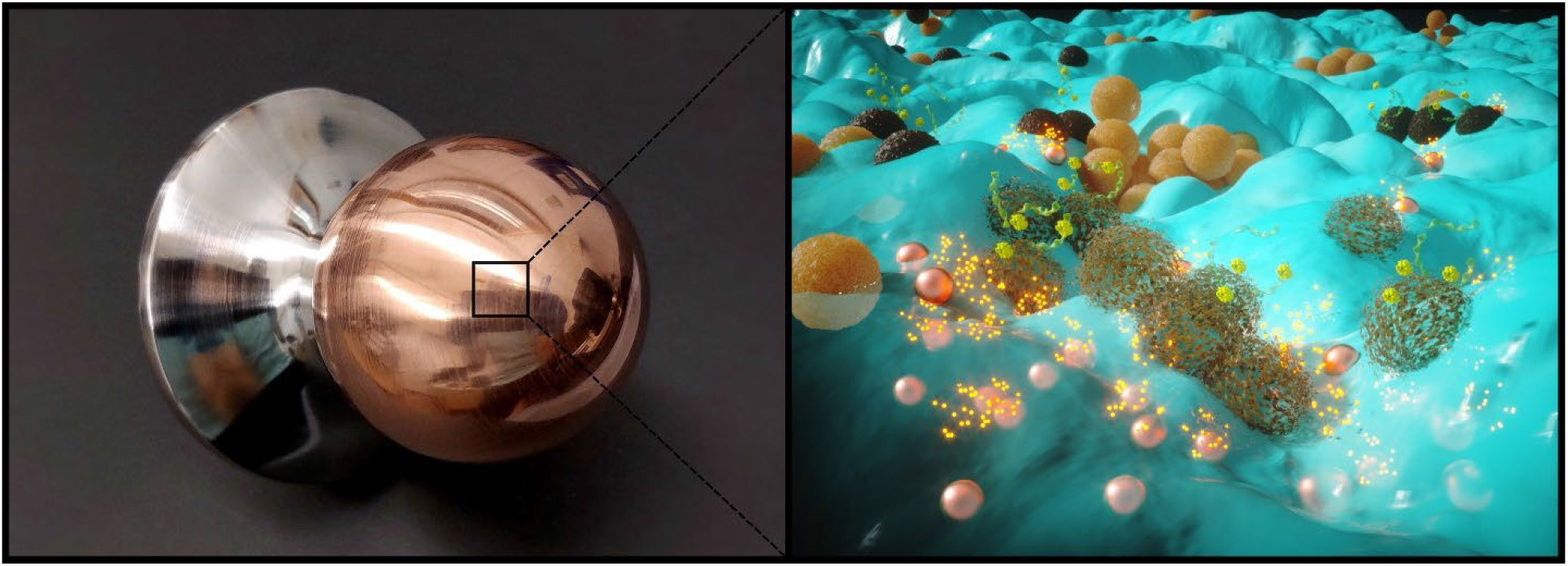

## 1. Introduction

High-touch surfaces (e.g., railings and doorknobs) in commercial, residential, and public infrastructure are potential reservoirs of pathogenic bacteria and viruses.^[1,2]^ The global SARS-CoV-2 pandemic has increased public awareness of the ability of high-touch surfaces to transmit infection.^[3,4]^ Cleaning and disinfection to mitigate the bioburden on hard surfaces is costly and cannot feasibly be conducted with sufficient frequency to be fully effective. Self-disinfecting surfaces made of or coated with copper (Cu) can minimize the spread of microorganisms and viruses on high-touch surfaces.^[5,6]^ At present, Cu is the only solid antimicrobial material registered as a self-sanitizer by Health Canada^[7]^ and the U.S. Environmental Protection Agency (EPA), and in 2021, it was confirmed that it is also antiviral, inactivating 99.9% of SARS-CoV-2 virus particles within two hours.^[8]^

Repeated touch as well as cleaning/disinfection of Cu-bearing surfaces triggers the release of Cu ions or molecular compounds containing Cu(I) or Cu(II) oxides through aqueous corrosion.^[9]^ It is well known that the bactericidal behavior of a Cu-based surface is mainly due to its capacity to release Cu ions.^[10–13]^ Thus, ensuring the rapid and continuous release of Cu ions from the Cu surface is critical to quickly killing pathogens, as well as maintaining Cu’s self-sanitizing properties long-term.^[12]^ Cu ions kill bacteria by denaturing the cell membrane^[14]^ and intracellular proteins, as well as altering the structure of DNA, and enhancing the production of destructive reactive oxygen species (ROS).^[15]^

Despite the proven efficacy of Cu as an antimicrobial agent, the bactericidal activity of Cu proceeds relatively slowly (e.g., 2 h for 98% reduction in *Staphylococcus aureus*).^[11]^ Further, in healthcare settings, Cu appears to be less effective against Gram-positive bacteria than Gram-negative bacteria.^[16,17]^ This difference is attributed to the variation in peptidoglycan wall thickness,^[13,17]^ which is approximately an order of a magnitude thicker for Gram-positive bacteria compared to Gram-negative.^[18]^ The Gram-positive bacterium, *S. aureus*, is responsible for skin and wound infections and is increasingly resistant to antibiotics. According to the World Health Organization, antibiotic resistance is a major threat to global health and food safety.^[19]^ It leads to longer hospitalizations, higher medical costs, and higher mortality rates. Nosocomial infections cause considerable patient morbidity and mortality.^[20,21]^ As such, developing new technology to mitigate their spread is critical.

In this study, we sought to mitigate surface-based transmission of infectious diseases through development of a multifunctional Cu coating, engineered to enhance its antibacterial efficacy towards Gram-positive bacteria. Our coating technology combines Cu nanoparticles with a cationic polymer, linear polyethyleneimine (LPEI), which has demonstrated antimicrobial activity against pathogenic Gram-positive and -negative bacteria as well as viruses.^[15,22]^ In addition, our nanocomposite copper coatings contain hierarchical surface features on both the micro- and nanoscale, that increases their hydrophobicity and thus hinders the ability of bacteria to adhere to the surface. The incorporation of nanotopography has also been found to kill bacteria by rupturing their cell wall,^[23]^ as has been observed in some species of dragonflies and geckos with nanostructured skins.^[24]^ Finally, by leveraging the difference in standard reduction potential between Cu (0.34 V vs. SHE) and Zn (-0.76 V vs. SHE), we were able to trigger the rapid release of biocidal metal ions from the coating, rather than waiting for Cu ions to leach via aqueous corrosion.

## 2. Results and Discussion

### 2.1 Coating Morphology and Chemistry

Nanocomposite Cu coatings with nanoscale or multiscale (micro- and nanoscale) surface features were fabricated by tuning the EPD processing parameters. When glacial acetic acid is added to a mixture containing LPEI in H_2_O, LPEI dissolves and becomes protonated, forming a charged suspension:

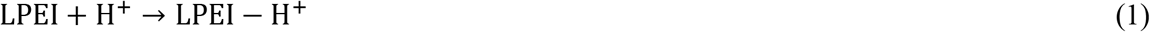

LPEI-H^+^ then acts as a charging and dispersing agent for Cu nanoparticles or metal salts (CuCl_2_ or ZnCl_2_) added to the suspension, by adsorbing to the Cu nanoparticle surface or metal cation via the amine group in the LPEI monomer. The mechanism of deposition of the LPEI-Cu nanocomposite coating is based on 1) cataphoresis of the LPEI-Cu nanoparticle complex, 2) cathodic electrosynthesis of Cu oxide and Cu hydroxide, and 3) charge neutralization.^[25]^

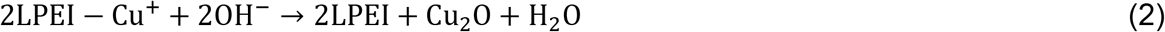

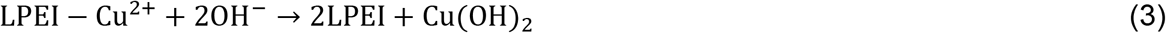

To determine the optimal processing parameters for obtaining nano- or multiscale surface structuring, we studied the influence of suspension pH and electrodeposition potential on contact angle (see Fig. SI1, Supporting Information), and surface morphology. Based on these results, we determined that the optimal pH and deposition potential were 5.3 and 40 V, respectively. Moreover, we discovered that the mean contact angle of the Cu nanocoatings could be increased from 83° to 110° (Fig. 1), through the addition of metal salts, CuCl_2_ and ZnCl_2_, to the colloidal precursor. This is most likely due to the introduction of cauliflower-like microscale surface features (Fig. 1b-c) and increased surface roughness, which has been shown to change the wetting behaviour of surfaces.^[26]^ This effect can be explained using DLVO (Derjaguin–Landau–Verwey–Overbeek) theory, which considers the forces between charged particles interacting through a suspension.^[27]^ Using EPD, the final coating morphology can be tailored by tuning the colloidal stability of the precursor suspension. Specifically, smooth, uniform, coatings are fabricated from stable colloidal suspensions, while coatings with nano- or hierarchical surface features are deposited from quasi-stable suspensions, by depositing multiple layers of randomly oriented coagulated particles.^[28]^

**Figure 1.**
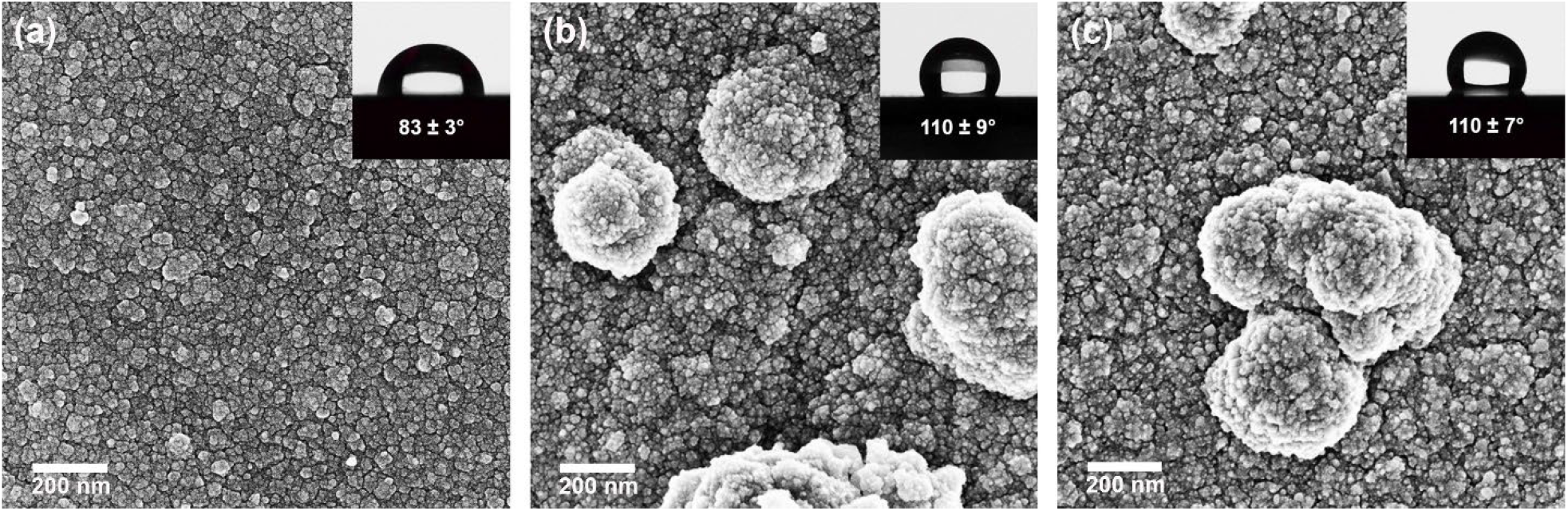
Scanning electron micrographs and mean (± SD; *n* = 5) contact angles of nanocomposite Cu coatings a) NP1 (0 mg L^-1^ metal salt), b) NP2 (100 mg L^-1^ CuCl_2_), and c) NP3 (500 mg L^-1^ ZnCl_2_).

The colloidal suspension used to fabricate NP1 was relatively stable, compared to colloidal precursors used to fabricate NP2 and NP3, which was reflected in the final coating morphology, which contained uniformly distributed nanoscale surface features (Fig. 1a). The addition of CuCl_2_ and ZnCl_2_ increased the ionic concentration of the suspension, which decreased the thickness of the electrical double-layer. As mentioned earlier, this causes the contribution from electrostatic repulsion to become less dominant than the attractive London-van der Waals forces, which decreases the overall stability of the suspension.^[29]^ However, the metal salt concentrations were not high enough to make the suspension unstable and cause particle flocculation. Rather, the suspension became quasi-stable, which resulted in the formation of irregular deposition features (i.e., cauliflower features in Fig. 1b and 1c) and therefore hydrophobicity of the NP2 and NP3 coatings.

Although NP1 had a lower contact angle than NP2 and NP3, the contact angle for all nanocomposite Cu coatings were higher than PCu (53 ±2°; see Fig. SI2, Supporting Information). Moreover, they exhibited strong coating adhesion to the SS substrate, with all coatings obtaining 4B classification (<5% detachment) according to the ASTM D-3359-17 (Tape Test) (see Fig. SI3 in the Supporting Information).

XPS revealed the surface of all nanocomposite Cu coatings consisted primarily of Cu_2_O and Cu(OH)_2_ (see Fig. 2), likely due to a combination of electrosynthesis of Cu oxide and hydroxide during deposition, as well as atmospheric oxidation of Cu nanoparticles prior to deposition. However, the XPS spectra showed that these coatings also contained Cu(0), due to deposition of pure metallic Cu nanoparticles and the direct reduction of Cu(I) and Cu(II) ions from the suspension. ToF-SIMS analysis of the nanocomposite Cu coatings confirmed the presence of a CuCl_2_^-^ complex in NP3 coatings (see Fig. SI4 in the Supporting Information) before and after bactericidal efficacy evaluations. This complex probably forms because it is more stable than ZnCl_2_, because CuCl_2_^-^ has a higher cumulative stability constant than ZnCl_2_.^[30,31]^

**Figure 2.**
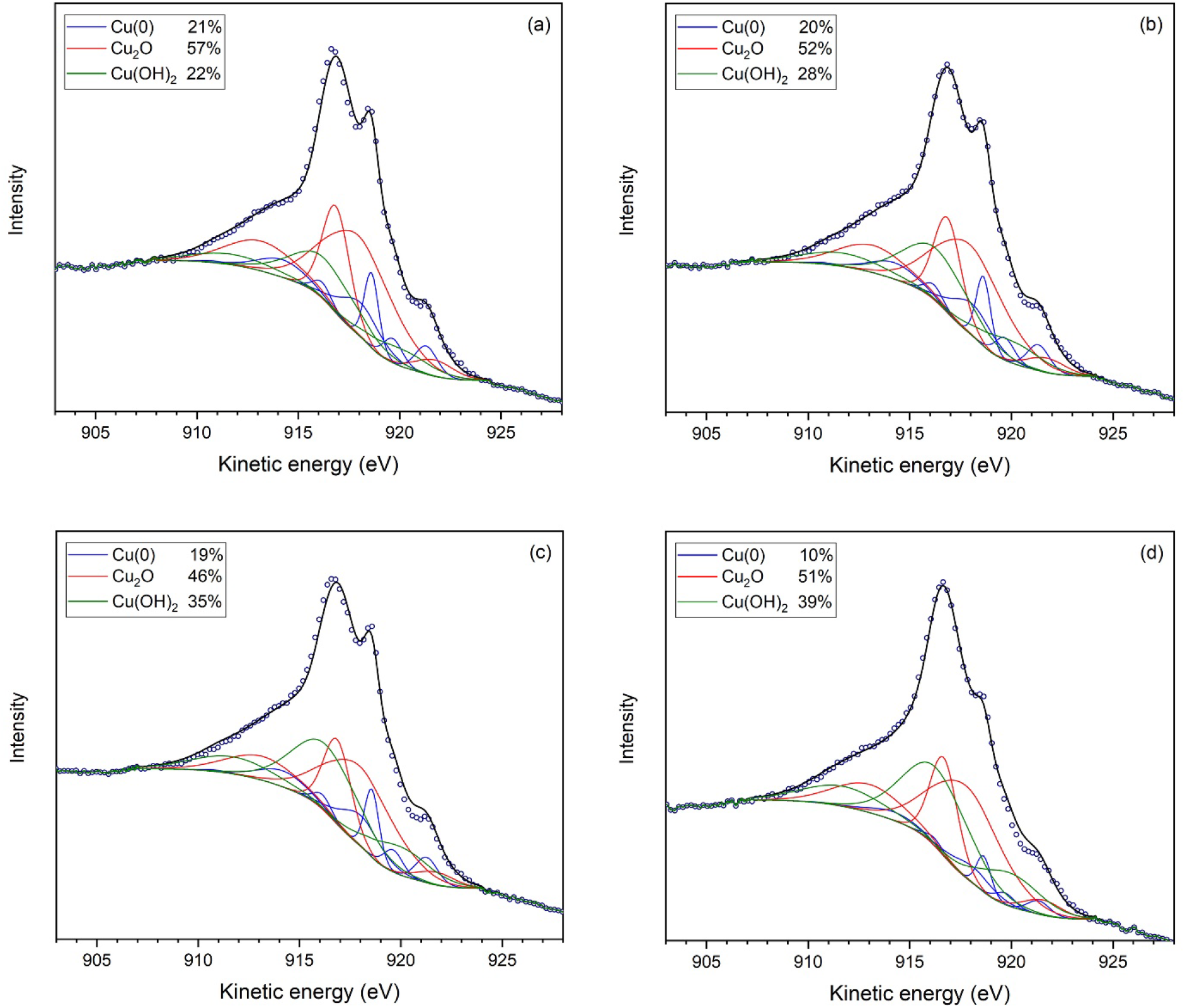
Deconvoluted Cu LMM Auger spectra and relative fraction of Cu species of a) pure Cu and nanocomposite Cu coatings b) NP1 (0 mg L^-1^ metal salt), c) NP2 (100 mg L^-1^ CuCl_2_), and d) NP3 (500 mg L^-1^ ZnCl_2_).

XPS and ToF-SIMS analyses did not detect Zn on the surface of NP3, likely because the analysis depths of these techniques are shallow (<10 nm and 1 nm, respectively).^[32],[33]^ In addition, selective leaching of Zn in the presence of Cu is spontaneous and highly-favourable, since it has a stronger tendency to oxidize than Cu,^[34]^ which also could contribute to our inability to detect Zn using XPS and ToF-SIMS. In contrast, ICP-OES showed progressive Zn ion release from NP3 (Fig. 3b), confirming its presence in the coating. Therefore, it is likely that the topmost layer of the deposited film was free of Zn— as XPS and ToF-SIMS analysis suggest—and Zn was present in the underlying layer.

**Figure 3.**
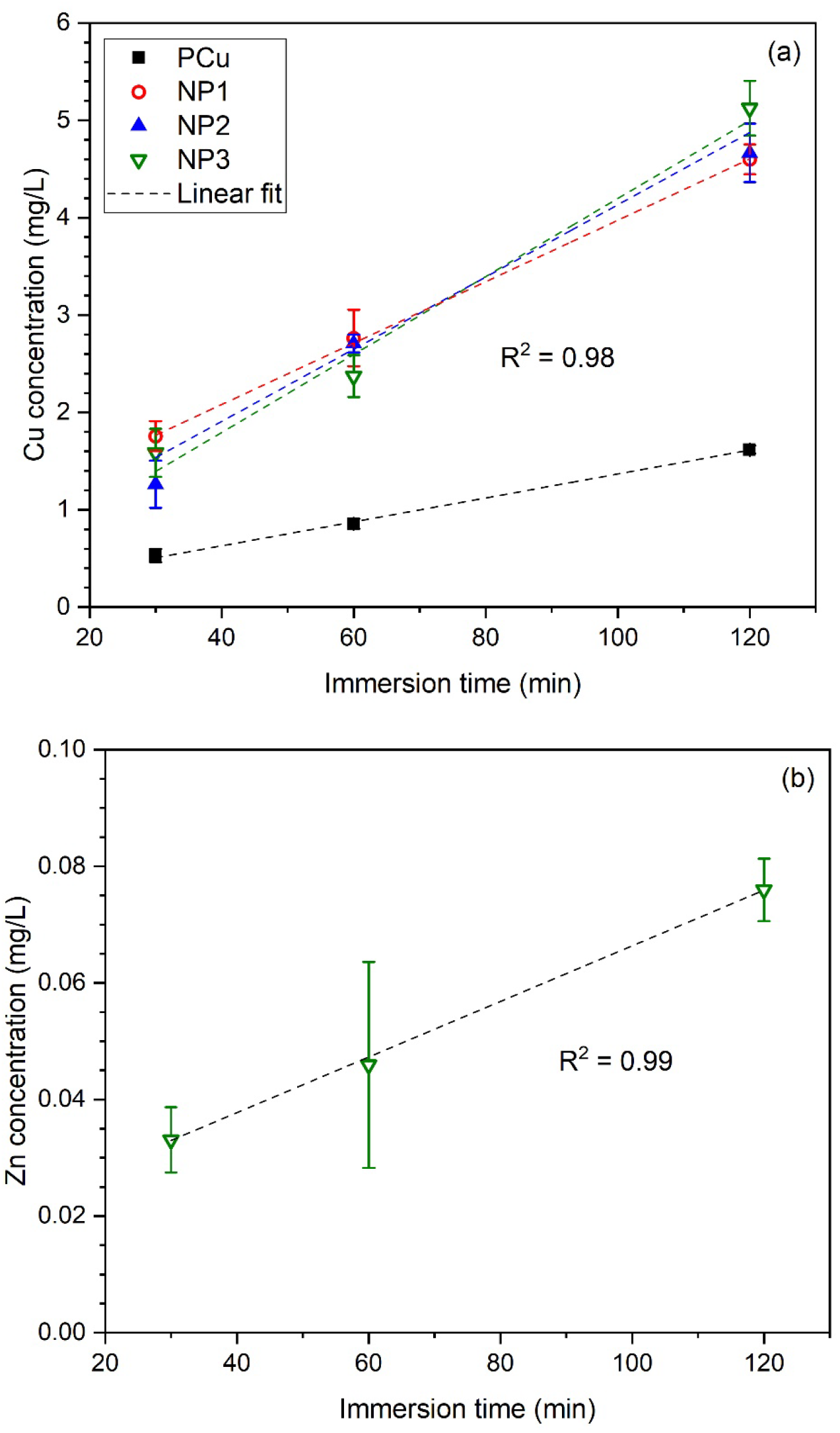
Linear regression of mean (±SD, *n* = 3) a) Cu concentration vs. immersion time in phosphate buffer solution for pure Cu (PCu) and Cu-nanocoated stainless steel coupons NP1 (0 mg L^-1^ metal salt), NP2 (100 mg L^-1^ CuCl_2_), and NP3 (500 mg L^-1^ ZnCl_2_); b) Zn concentration vs. immersion time in phosphate buffer solution for Cu-nanocoated stainless steel coupons NP3.

The fraction of Cu(0) present on the surface of PCu (Fig. 2a) and nanocomposite Cu coatings NP1 and NP2 (Fig. 2b–c) was similar (~20%) and higher than the NP3 coating (10%; Fig. 2d). All three nanocomposite coatings were enriched with Cu(OH)_2_ relative to PCu, especially NP3. The thickness of the native oxide film on pure Cu is 3–4 nm,^[35,36]^ and the effective analysis depth of XPS is <10 nm;^[32]^ therefore, some of the Cu(0) signals for PCu likely originated from the underlying bulk Cu. Zn was detected on the NP3 surface in trace concentrations, which made the signal-to-noise ratio too small for accurate fitting. XPS survey spectra for as-polished PCu and as-deposited nanocomposite Cu coatings are shown in Fig. SI5 in the Supporting Information.

### 2.2 Cu Ion Release

As the antibacterial properties of Cu and its efficacy are dependent on the rate of Cu ion release, we compared the Cu release rate from our nanocomposite Cu coatings to PCu in PBS. Cu ion concentrations increased linearly with increasing immersion time for all samples (*R*^2^ = 0.98; Fig. 3). However, PCu released Cu at a much lower rate than nanocomposite Cu coatings NP1–3, likely due to the passivation of the PCu surface, which impeded anodic dissolution.^[37]^ Although the low signal-to-noise ratio prevented accurate fitting during XPS analysis, Zn ion release was detected with ICP-OES analysis, as noted earlier. Zn ion concentration also increased linearly with immersion time for nanocoating NP3 (Fig. 3b).

### 2.3 Antibacterial Efficacy

Compared to SS, PCu and SS coated with NP1–3 had a 95.1–99.9% reduction in viable *S. aureus* CFUs (Table 2). The mean amount remaining on disk surfaces after swabbing with Quick Swabs was 0.2%. Compared to PCu and for a given post-inoculation recovery time, NP3 had the highest percent reduction in viable *S. aureus* CFUs such that at 120 minutes post-inoculation, it was the only nanocoating to meet U.S. Environmental Protection Agency standards as a sanitizer.

**Table 1:** Experimental design for ion release and bactericidal experiment with pure copper coupons and nanocoated (linear polyethyleneimine + Cu nanoparticles) stainless steel coupons (NP1–NP3).

| Coupon | Phosphate Buffer Saline (PBS) Soak Time/Bacteria Recovery Time (min) |  |  |
| --- | --- | --- | --- |
|  | 30 | 60 | 120 |
|  | Replicates |  |  |
| Pure copper | 3 | 3 | 3 |
| NP1 (no metal salt) | 3 | 3 | 3 |
| NP2 (100 mg L<sup>-1</sup> CuCl<sub>2</sub>) | 3 | 3 | 3 |
| NP3 (500 mg L<sup>-1</sup> ZnCl<sub>2</sub>) | 3 | 3 | 3 |

**Table 2:**
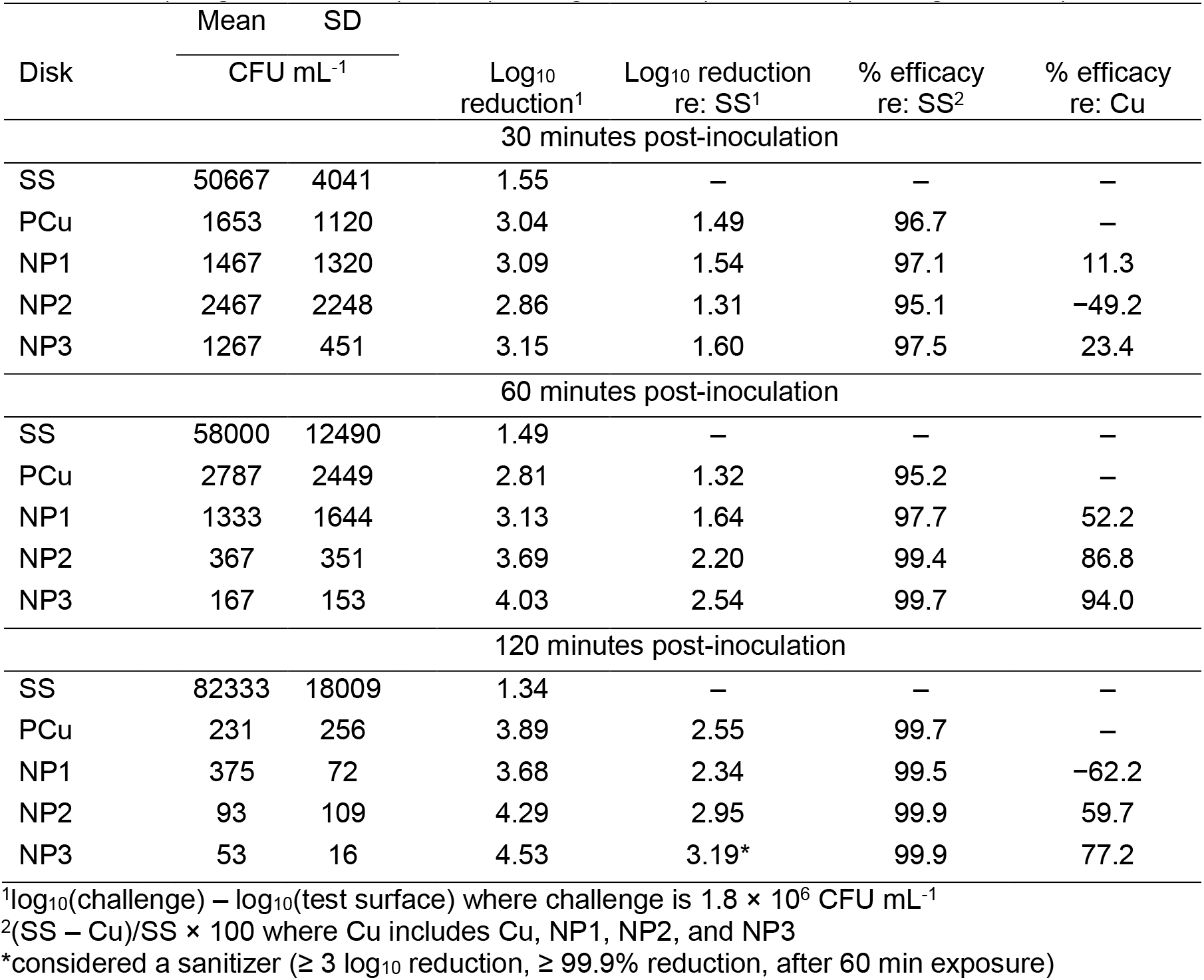
Bactericidal activity against *S. aureus* for stainless steel (SS), pure Cu (PCu), and SS disks coated with NP1 (0 mg L^-1^ metal salt), NP2 (100 mg L^-1^ CuCl_2_), and NP3 (500 mg L^-1^ ZnCl_2_)

*P. aeruginosa* samples were too dilute at 30 and 60 min to determine optimal bactericidal activity of the Cu nanocoatings (Table 3). Among the colonies that could be enumerated, there was a trend for the increased bactericidal activity of the PCu and NP3 products at 60 minutes post-inoculation compared to NP1 and NP2 such that they met the U.S. Environmental Protection Agency standards as sanitizers. At 120 minutes post-inoculation, PCu and NP1–3 all met U.S. Environmental Protection Agency standards as sanitizers.

**Table 3:** Bactericidal activity against *P aeruginosa* for stainless steel (SS), pure Cu (PCu), and SS disks coated with NP1 (0 mg L^-1^ metal salt), NP2 (100 mg L^-1^ CuCl_2_), and NP3 (500 mg L^-1^ ZnCl_2_)

| Disk | Mean | SD | Log<sub>10</sub> reduction<sup>1</sup> | Log<sub>10</sub> reduction re: SS<sup>1</sup> | % efficacy re: SS<sup>2</sup> | % efficacy re: Cu |
| --- | --- | --- | --- | --- | --- | --- |
|  | CFU mL<sup>-1</sup> |  |  |  |  |  |
| 30 minutes post-inoculation |  |  |  |  |  |  |
| SS | 12667 | 4041 | 2.07 | – | – | – |
| PCu | 67 | 115 | 4.35 | 2.28 | 99.5 | – |
| NP1 | 0** | 0** | 6.18 | 4.10* | 100.0 | 100.0 |
| NP2 | 33 | 58 | 4.66 | 2.58 | 99.7 | 50.7 |
| NP3 | 0** | 0** | 6.18 | 4.10* | 100.0 | 100.0 |
| 60 minutes post-inoculation |  |  |  |  |  |  |
| SS | 3333 | 1528 | 2.65 | – | – | – |
| PCu | 0** | 0** | 6.18 | 3.52* | 100.0 | – |
| NP1 | 33 | 58 | 4.66 | 2.00 | 99.0 | – |
| NP2 | 133 | 231 | 4.05 | 1.40 | 96.0 | – |
| NP3 | 0** | 0** | 6.18 | 3.52* | 100.0 | – |
| 120 minutes post-inoculation |  |  |  |  |  |  |
| SS | 11667 | 4041 | 2.11 | – | – | – |
| PCu | 1 | 2 | 6.18 | 4.07* | 100.0 | – |
| NP1 | 0** | 1 | 6.18 | 4.07* | 100.0 | 100.0 |
| NP2 | 0** | 0** | 6.18 | 4.07* | 100.0 | 100.0 |
| NP3 | 0** | 1 | 6.18 | 4.07* | 100.0 | 100.0 |

<sup>1</sup>log<sub>10</sub>(challenge) – log<sub>10</sub>(test surface) where challenge is 1.5 × 10<sup>6</sup> CFU mL<sup>-1</sup>
<sup>2</sup>(SS – Cu)/SS × 100 where Cu includes Cu, NP1, NP2, and NP3
\*considered a sanitizer (≥ 3 log<sub>10</sub> reduction, ≥ 99.9% reduction, after 60 min exposure)
\*\* log<sub>10</sub>(test surface) was assumed to be equal to zero for surfaces when the bacteria remaining on the surface was less than 0.5 CFU mL<sup>-1</sup>

At 30 minutes post-inoculation, MALDI-TOF identified *Bacillus cereus* on the sonicated *S. aureus* and *P. aeruginosa* Cu blood agar plates. One other contaminant was noted on a blood agar plate inoculated with *S. aureus:* NP1 at 120 minutes post-inoculation. The colony could not be identified by MALDI.

During recovery of bacteria, slight discoloration or oxidation of the Cu was observed on NP1–3 nanocoated disks. This was absent from SS disks and partially visible on PCu disks. XPS analysis of the Cu nanocoatings after bactericidal efficacy evaluations revealed that disk surfaces were mainly covered with Cu_3_(PO_4_)_2_ and with some of Cu(OH)_2_ and Cu_2_O. PCu coupons, however, were entirely covered with Cu(OH)_2_. High-resolution XPS spectra obtained from the surface of the nanocomposite Cu coatings and PCu after bactericidal efficacy evaluations can be found in Fig. SI6 in the Supporting Information.

To evaluate the bacterial viability of *S. aureus* and *P. aeruginosa* on PCu compared to NP3, we used live/dead cell staining. Live cells were stained using SYTO™ 9, which enters all cells, binds to DNA and RNA, and emits green fluorescence. PI also binds to DNA and RNA, but can only enter cells with a compromised membrane and emit red fluorescence. As can be seen in Fig. 4, both types of bacteria were more viable on PCu compared to NP3. However, it should be noted that the morphology of all bacteria on Cu-containing surfaces was different compared to those on SS (See Fig. SI7, Supporting Information), indicating cellular damage.

**Figure 4.**
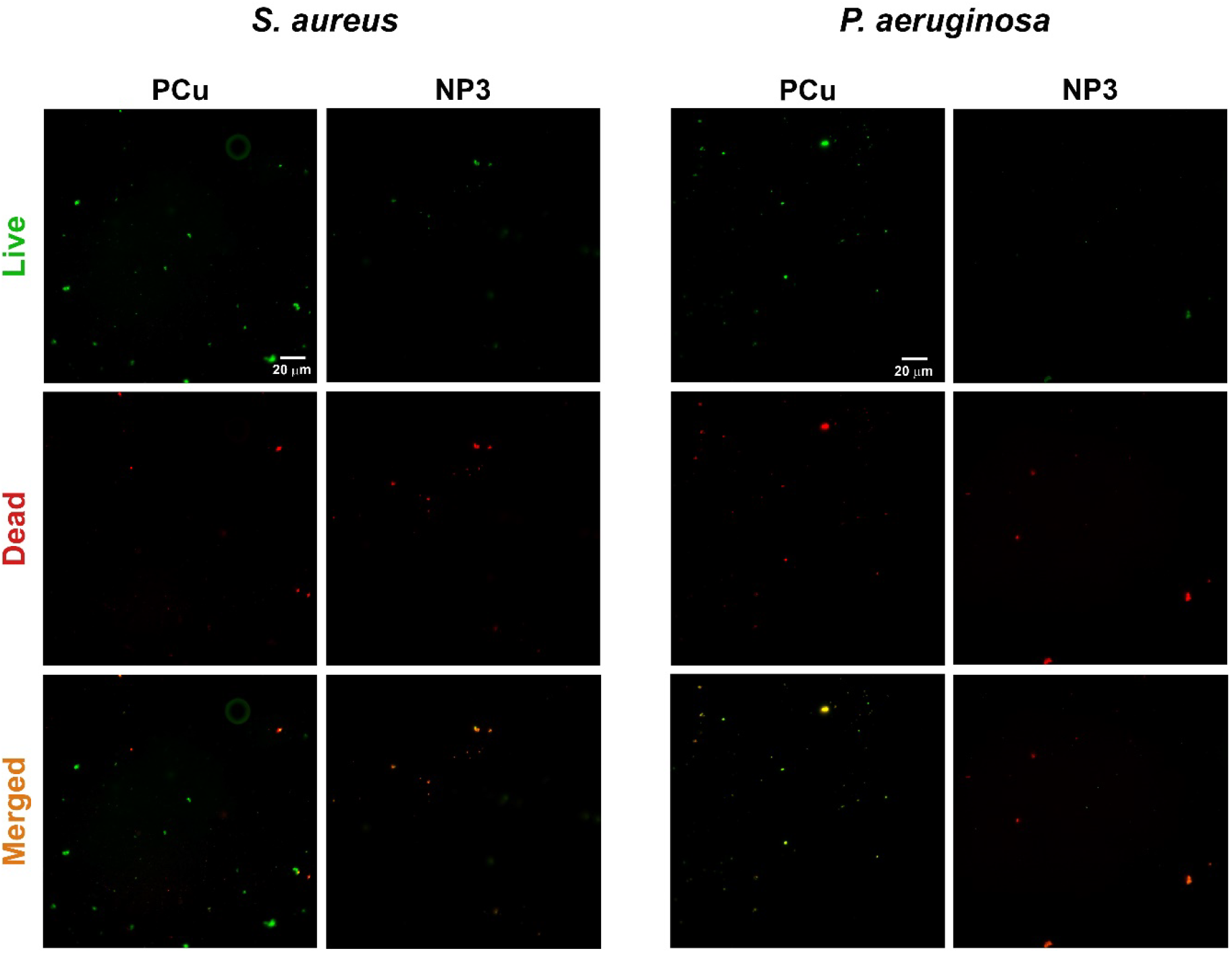
Bacterial viability of *S. aureus* and *P. aeruginosa* on PCu and NP3. The live/dead bacterial cells were stained at 60 min by live/dead assay, and representative images after cell staining were observed using a fluorescent microscope.

A possible explanation for the superior bactericidal efficacy of NP3 is that the presence of Zn in the NP3 coating drives the rapid and selective leaching of Zn ions, due to its lower standard reduction potential compared to Cu. Similar to Cu, Zn is also biocidal, and the toxicity of ZnO nanoparticles to Gram-positive and -negative bacteria and viruses is well-studied.^[38],[39]^ When bacteria are exposed to ZnO nanoparticles, biocidal Zn^2+^ ions are released, which destroy the bacteria cell wall and generates ROS.^[38,40–42]^ In addition, NP3 coatings had the least amount of metallic Cu. This suggests that Cu ions, either as Cu(I) or Cu(II), were readily available to release from the surface to kill bacteria. In contrast, a higher fraction of metallic Cu on PCu, NP1, and NP2 implies that these surfaces require oxidation of Cu(0) (i.e., anodic dissolution of Cu(0)) prior to releasing Cu ions, which is another possible mechanism for higher efficacy of NP3 coating compared to PCu, NP1, and NP2.

## 3. Conclusions

Nanostructured Cu coatings comprising Cu nanoparticles and LPEI were developed and applied on a SS substrate using EPD. The addition of CuCl_2_ and ZnCl_2_ salts to the coatings led to the formation of surface roughness across multiple length scales, through the introduction of microscale cauliflower-shaped features, embedded within the existing nanostructured matrix. This increased coating hydrophobicity, which has been found to impact bacterial adhesion to surfaces. Several mechanisms are involved in the superior bactericidal efficacy of the ZnCl_2_-containing coating. First, the nanostructured surface features can stretch the cell envelope of adhered bacteria, which eventually leads to the tearing and subsequent leakage of essential intracellular materials for the survival of the bacteria. Moreover, compared to the as-polished PCu, the Cu nanocoatings have a higher specific surface area, which results in higher Cu ion release. Finally, adding Zn, a metal with a lower standard reduction potential than Cu, drives the rapid and selective leaching of antimicrobial Zn ions, which improved antibacterial efficacy.

## 4. Experimental Procedure

### 4.1 Materials and Reagents

LPEI (molecular weight 250,000) was purchased from Polysciences Inc. (Warrington, PA, USA). Glacial acetic acid, Cu nanoparticles (60–80 nm; ≥99.5% trace metals basis), phosphate-buffered saline (PBS), yeast extract, and mucin (from bovine submaxillary glands) were purchased from Sigma-Aldrich (Oakville, ON, Canada). Anhydrous ethanol (ACS Reagent Grade), CuCl_2_ (ACS Reagent Grade), ZnCl_2_ (ACS Reagent Grade), SYTO^TM^ 9 Green Fluorescent Nucleic Acid Stain, Propidium Iodide (1.0 mg mL^-1^ Solution in Water), ProLong™ Live Antifade Reagent (for live cell imaging), were purchased from ThermoFisher Scientific (Canada). 95% Ethyl Alcohol was purchased from Greenfield Global (Irvine, CA, USA). Bovine serum albumin (BSA) was purchased from Proliant Biologicals (Ankeny, IA, USA). Commercially pure copper (PCu) (99.90% grade C11000) and AISI 316L stainless-steel (SS) were purchased from McMaster-Carr (Elmhurst, IL, USA). Letheen broth was purchased from 3M™ (London, ON, Canada) and Remel Inc. (San Diego, CA, USA). Petrifilm™ aerobic count plates and Quick Swabs were also purchased from 3M™ (London, ON, Canada). 5% Sheep’s blood agar plates were purchased from Oxoid (Nepean, ON, Canada).

### 4.2 Coating Fabrication

Electrophoretic deposition (EPD) was used to fabricate the nanocomposite Cu coatings. Coatings were deposited from a colloidal suspension containing LPEI dissolved in a 20% water and 80% ethanol mixture (by volume) and Cu NPs. The suspension was prepared by first protonating 2 g L^-1^ LPEI with 0.5 mol L^-1^ glacial acetic acid in deionized water. While stirring, anhydrous ethanol was added to the suspension, to minimize H2 gas evolution at the cathode during deposition and improve coating quality. Cu nanoparticles were added to this mixture for a final concentration 1 g L^-1^, and the colloidal suspension was sonicated for 10 min. Either 100 mg L^-1^ CuCl_2_ or 500 mg L^-1^ ZnCl_2_ were subsequently added to the colloid to increase conductivity, modify the coating morphology, and in the case of ZnCl_2_-containing coatings, triggering the rapid release of biocidal metal ions. The three treatments are designated as NP1 (0 mg L^-1^ metal salt), NP2 (100 mg L^-1^ CuCl_2_), and NP3 (500 mg L^-1^ ZnCl_2_). Prior to EPD, the pH of all colloidal suspensions was adjusted to 5.3 using glacial acetic acid.

Coatings were deposited using a two-electrode electrochemical cell, SS cathode, and platinum anode. The surface area of both electrodes was 5 cm^2^, and the deposition potential was 40 V. The interelectrode distance was 15 mm, and the deposition time was 5 min at a temperature of 23 ± 1 °C.

### 4.3 Coating Characterization

The surface morphology of the nanocomposite Cu coatings was examined using field-emission scanning electron microscopy (ZEISS Sigma), and their contact angle as a function of deposition potential and suspension pH was determined using an Ossila goniometer. For each of five replicates, a 1–3 μL DI water droplet was dispensed on the surface, immediately illuminated, and photographed using the digital camera attached. The deposition conditions that yielded the highest contact angles were used for further material characterization and assays to evaluate antibacterial efficacy. Coating adhesion to the SS substrate was evaluated using the ASTM D3359-1720 tape-test technique.

Cu and Zn ion release from PCu coupons (surface area 5 cm^2^) and Cu-coated SS coupons was measured using inductively coupled plasma optical emission spectrometry (ICP-OES, Perkin Elmer, Optima 8300). The PCu coupons were wet ground with a 600 grit SiC emery paper and rinsed with DI water. The PCu and Cu-coated SS coupons were placed in beakers containing 15 mL phosphate-buffered saline for 30, 60, and 120 min at room temperature. Each condition was evaluated in triplicate (Table 1). Before ICP-OES, all samples were digested with concentrated HNO_3_ to dissolve suspended metals.

The surface chemical composition of PCu, NP1, NP2, and NP3 before and after antibacterial testing was determined using X-ray photoelectron spectroscopy (XPS) (see below). XPS can detect all elements except hydrogen and helium, probe the sample surface to a depth of 7–10 nm, and has a detection limit of 0.1–0.5 atomic %, depending on the element. XPS was carried out with a Kratos AXIS Supra X-ray photoelectron spectrometer (Surface Science Western, London, ON, Canada) using a monochromatic Al Kα source (15 mA, 15 kV). The instrument work function was calibrated to give a binding energy of 83.96 eV for the Au 4f7/2 line for metallic gold, and the spectrometer dispersion was adjusted to give a binding energy of 932.62 eV for the Cu 2p3/2 line of metallic copper. A Kratos charge neutralizer system was used on all specimens. Survey scan analyses were carried out with an analysis area of 300 × 700 μm and a pass energy of 160 eV. For high-resolution analyses, a pass energy of 20 eV was used. Spectra were charge-corrected to the mainline of the carbon 1s spectrum (adventitious carbon) set to 284.8 eV and were analyzed using CasaXPS software (ver. 2.3.23).

Determining the chemical state of Cu using XPS is challenging due to the complexity of the Cu 2p spectra resulting from overlapped binding energies for Cu(0) and Cu(I) species and transformation structures for Cu(II) species. Analysis of the Cu LMM Auger spectrum allows the various oxidation states of Cu to be quantified because it represents higher energy shifts compared to the Cu 2p.^[43]^ Cu species were deconvoluted according to the procedure outlined by Biesinger.^[43]^ The relative fraction of Cu species in each spectrum was calculated by the integration of peaks for different species.

Time-of-flight secondary ion mass spectroscopy (ToF-SIMS) was also used to assess the chemical composition of the NP1, NP2, and NP3 coating surfaces before and after antibacterial testing, using an ION-TOF (GmbH) (Surface Science Western, London, ON, Canada) equipped with a Bi cluster liquid metal ion source. A 25 keV Bi^3+^ cluster primary ion beam was pulsed at 10 kHz to bombard the sample surface and generate secondary ions. Positive or negative secondary ions were extracted from the sample surface, separated by mass, and detected via a reflectron-type TOF analyzer, allowing parallel detection of ion fragments having a mass/charge ratio up to ~900 within each cycle (100 μs). Each ion mass spectrum was calibrated using hydrogen and hydrocarbons, which are present on almost any surface. Ion mass spectra were collected using an area of 500 × 500 μm at 128 × 128 pixels.

### 4.4 Antibacterial Efficacy

Two days prior to the experiment, frozen stock of *S. aureus* ATCC 29213 or *P. aeruginosa* ATCC 27853 was streaked onto 5% sheep blood agar plates (BAP), and incubated for 18-24 hours at 37 °C. One day prior to the experiment, three representative colonies (different sizes) were selected from the overnight BAP, inoculated in 15 mL of tryptic soy broth (TSB) in a 200 mL flask, and incubated at 37 °C for 16 hours.

To evaluate the antibacterial efficacy of our nanocomposite copper coatings, SS disks with a diameter of 25mm and thickness of 3mm, were coated with NP1, NP2, or NP3 (Table 1) and compared to uncoated SS and PCu as control. All disks were cleaned in 95% ethanol and inoculated with a mixture of *S. aureus* ATCC 29213 or *P. aeruginosa* ATCC 27853 in simulated soil media (BSA, yeast, and mucin)^[44]^ to quantify bactericidal activity against Gram-positive and -negative bacteria, respectively. Bacteria were recovered from the disks at 30, 60, or 120 min after inoculation using 3M^TM^ Quick Swabs (Table 1). Undiluted,1:100 or 1:1000 dilutions of recovery broth were plated onto 3M^TM^ Petrifilm^TM^ Aerobic Count Plates and incubated for 2 days at 30 °C. Colony forming units were enumerated using a 3M^TM^ Petrifilm^TM^ Plate Reader Advanced system. This method can test five surfaces in triplicate at three time points within 3 h. Importantly, it allows for recovery of organisms at 30 min, which is the average drying time for the disks.

Swabbed coupons were then sonicated in Letheen broth, subcultured onto 5% sheep blood agar plates, and incubated for two days at 37 °C to assess the efficacy of the Quick swabs at bacterial retrieval and to assess contamination. The recovery broth was also subcultured to assess contamination, and Matrix-assisted laser desorption/ionization (MALDI-TOF) was used to identify aberrant colonies.

### 4.5 Live/Dead Staining

Bacterial suspensions were prepared using the same methods described above. Bacteria was inoculated onto the test material (SS, PCu, NP1, NP2, and NP3) and recovered after 30 or 60 minutes in 100 μL PBS. Recovered bacteria was immediately subjected to a staining cocktail containing equal parts of SYTO™ 9 and Propidium Iodide (PI) and a 1:100 diluted mixture of ProLong™ Live Antifade, for a final volume of 200 μL/ stain. Bacteria was incubated for 15-20 minutes in the staining cocktail in the dark, and then 10 μL was plated onto a labelled microslide for imaging. An Olympus BX61 optical microscope (BC Children’s Hospital Research Institute, Vancouver, BC, Canada) with 60X objective lens and an oil immersion was used to image bacteria on the PCu and nanocomposite Cu surfaces. A FITC filter was used to quantify live cells, since the emission maxima for SYTO™ 9 occurs at approximately 480/500 nm, while a TRITC was used to visualize dead cells, which has an emission maximum around 490/635 nm.

## Acknowledgements

The authors would like to gratefully acknowledge Teck Resources Limited for financially supporting this work.

## Data Availability Statement

The data that support the findings of this study are available from the corresponding author upon request.

