## Supporting Information for "An Engineered Nanocomposite Copper Coating with Enhanced Antibacterial Efficacy"

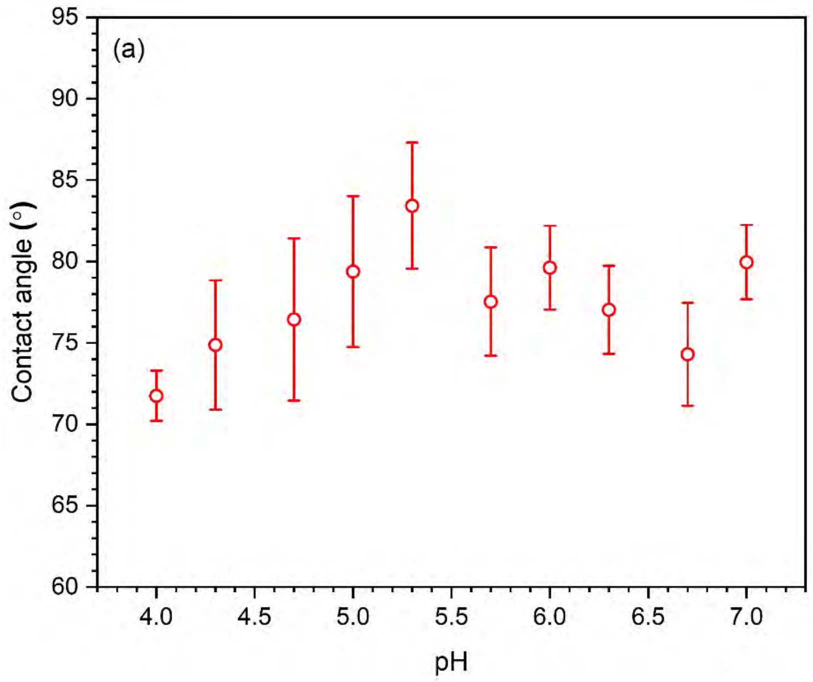


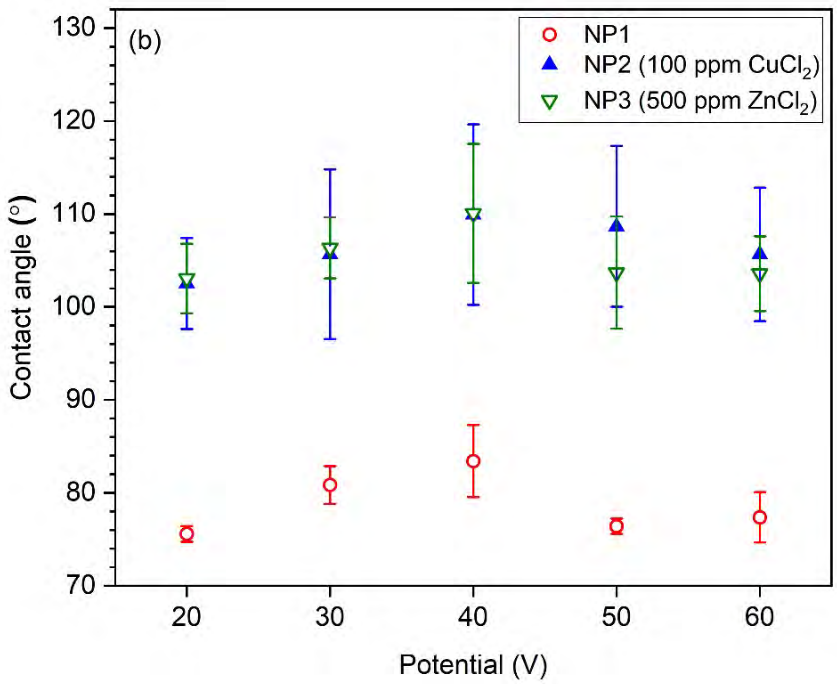


**Figure SI1.** Mean (± SD; $n$ = 5) contact angles for nanocomposite Cu coatings: a) NP1 coating at EPD potential of 40 V vs. pH, b) NP1, NP2, and NP3 coatings vs. EPD potential at pH 5.3. Addition of $CuCl_2$ and $ZnCl_2$ increased the contact angle through introducing dual-scale surface features by decreasing the stability of the colloidal precursor used for EPD.

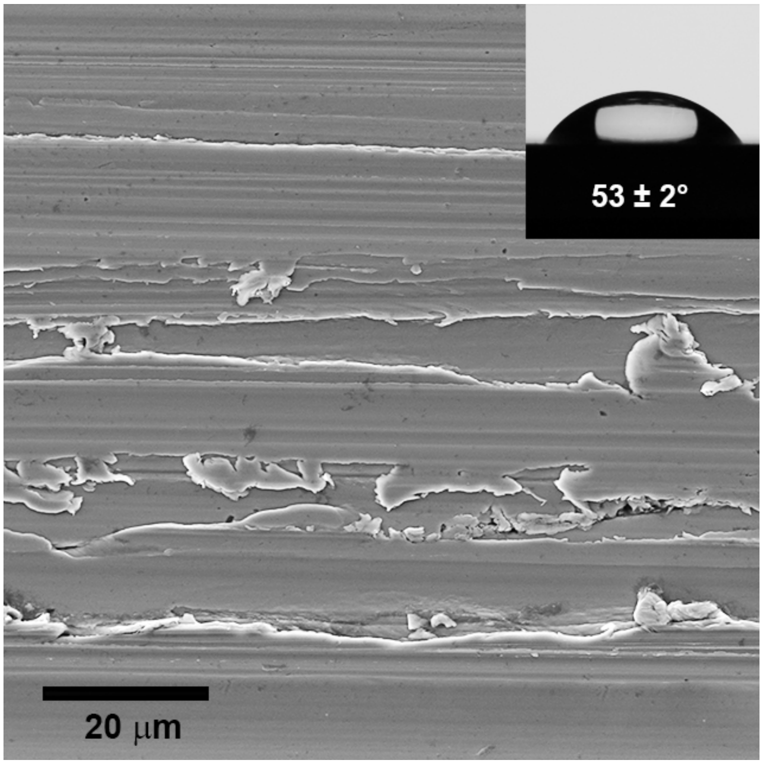


**Figure SI2.** Surface morphology and mean (± SD; $n$ = 5) contact angle of pure copper after wet grinding with a 600 grit SiC emery paper.

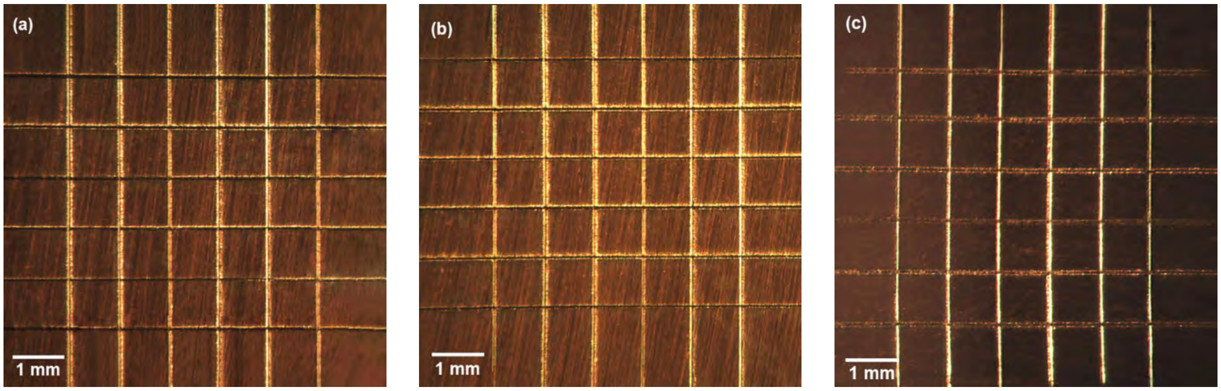


**Figure SI3.** Coating adhesion results for a) NP1 (0 mg $L^{-1}$ metal salt), b) NP2 (100 mg $L^{-1}$ $CuCl_2$), and c) NP3 (500 mg $L^{-1}$ $ZnCl_2$), determined using the ASTM D-3359-17 (Tape Test). All coatings exhibited 4B adhesion to the stainless-steel substrate.

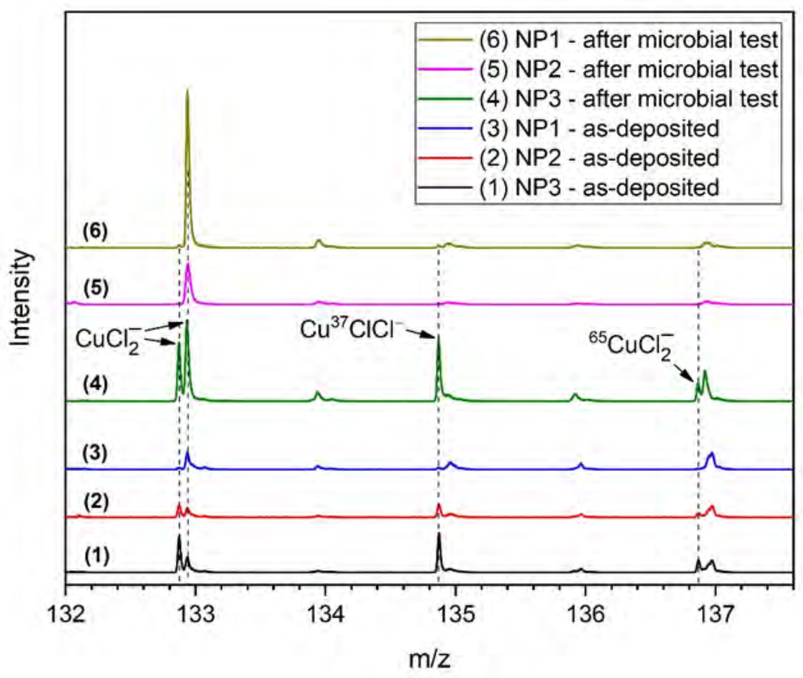


**Figure SI4.** Negative secondary ion mass spectra showing $CuCl_2^-$ in Cu nanocoating NP3

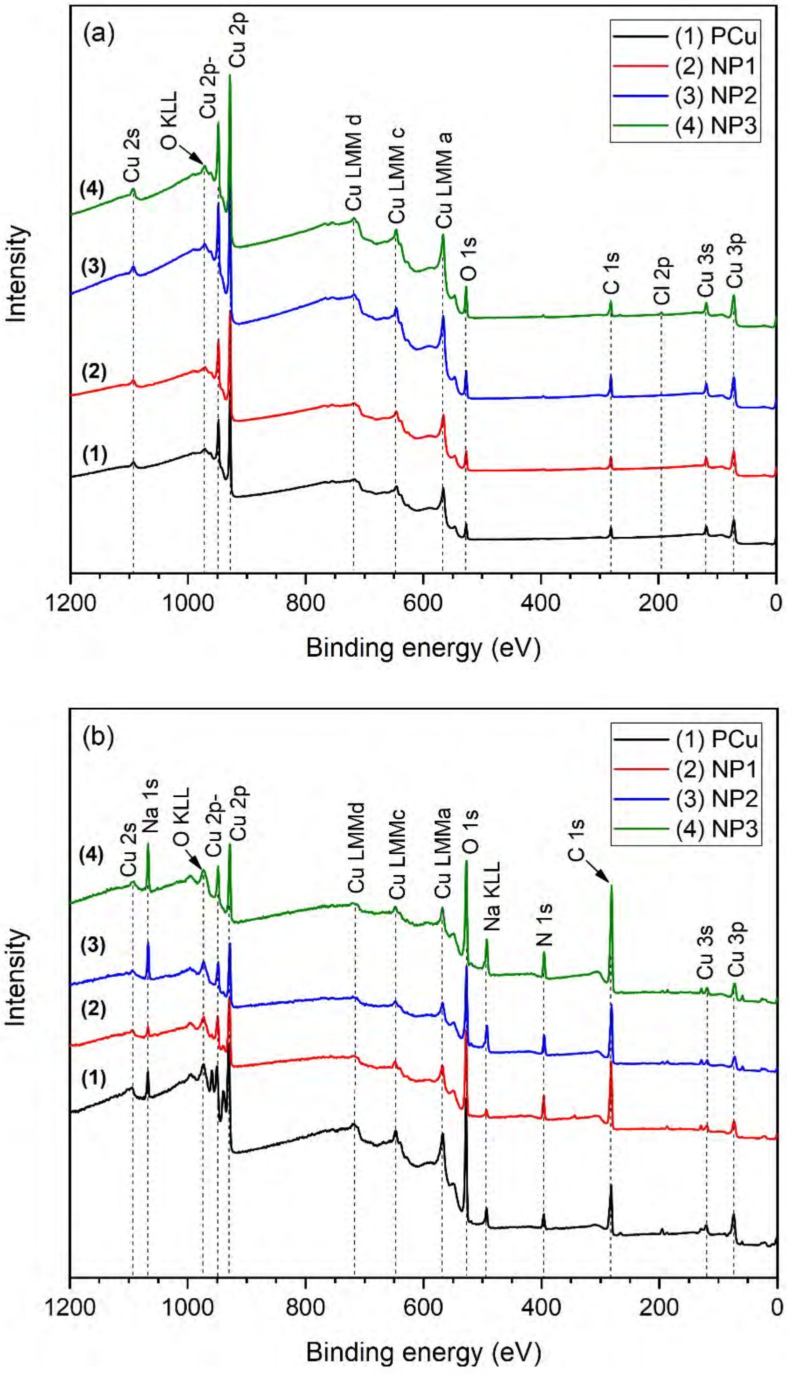


**Figure SI5.** X-ray photoelectron spectroscopy survey spectra for as-polished PCu and as-deposited nanocomposite Cu coatings NP1, NP2, and NP3 a) before and b) after bactericidal efficacy evaluations.

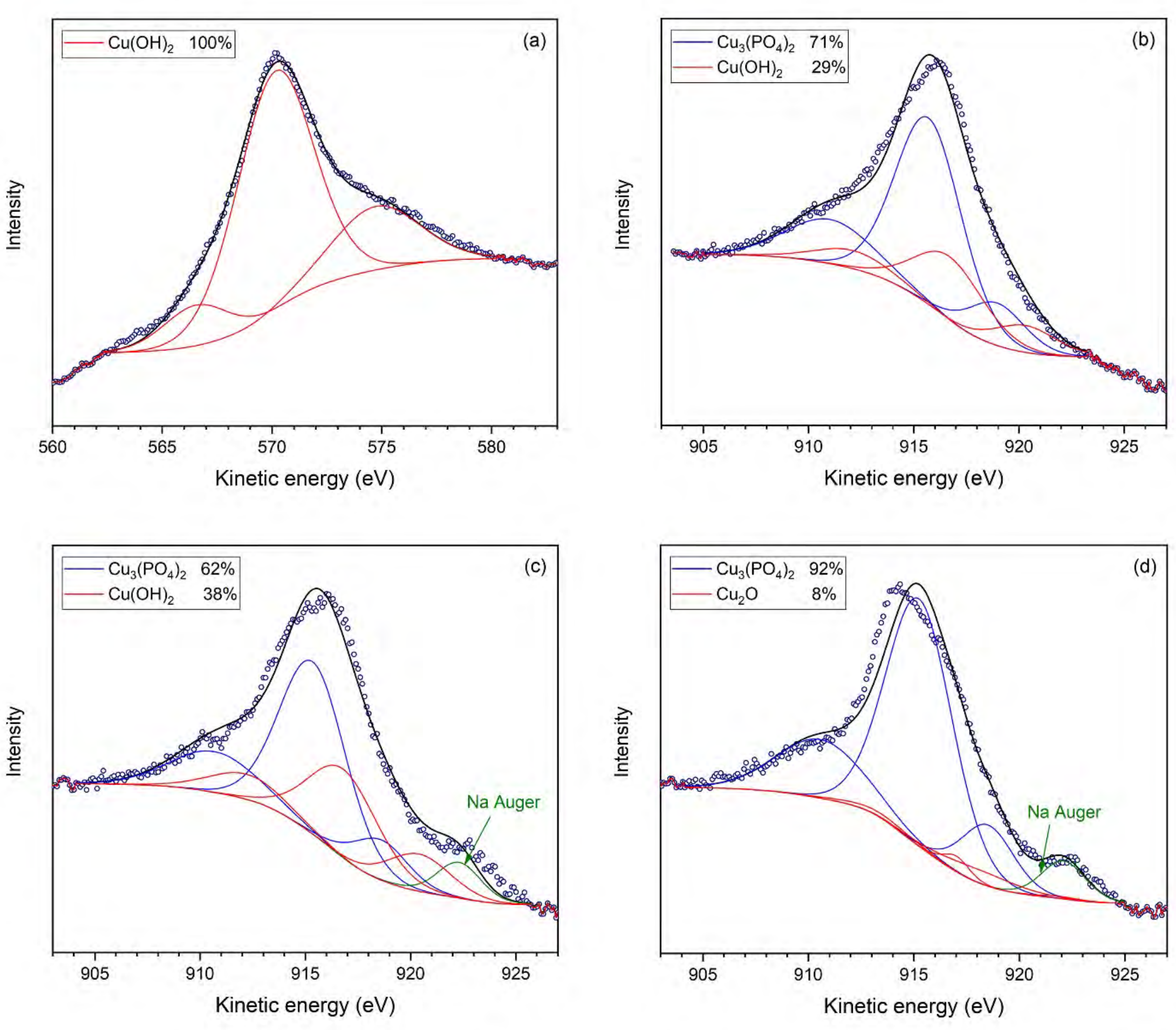


**Figure SI6.** Deconvoluted Cu LMM Auger spectrum after bactericidal efficacy evaluations of: a) PCu, b) NP1, c) NP2, and d) NP3. The relative fraction of Cu species was calculated using curve-fitting of the Cu LMM spectrum.

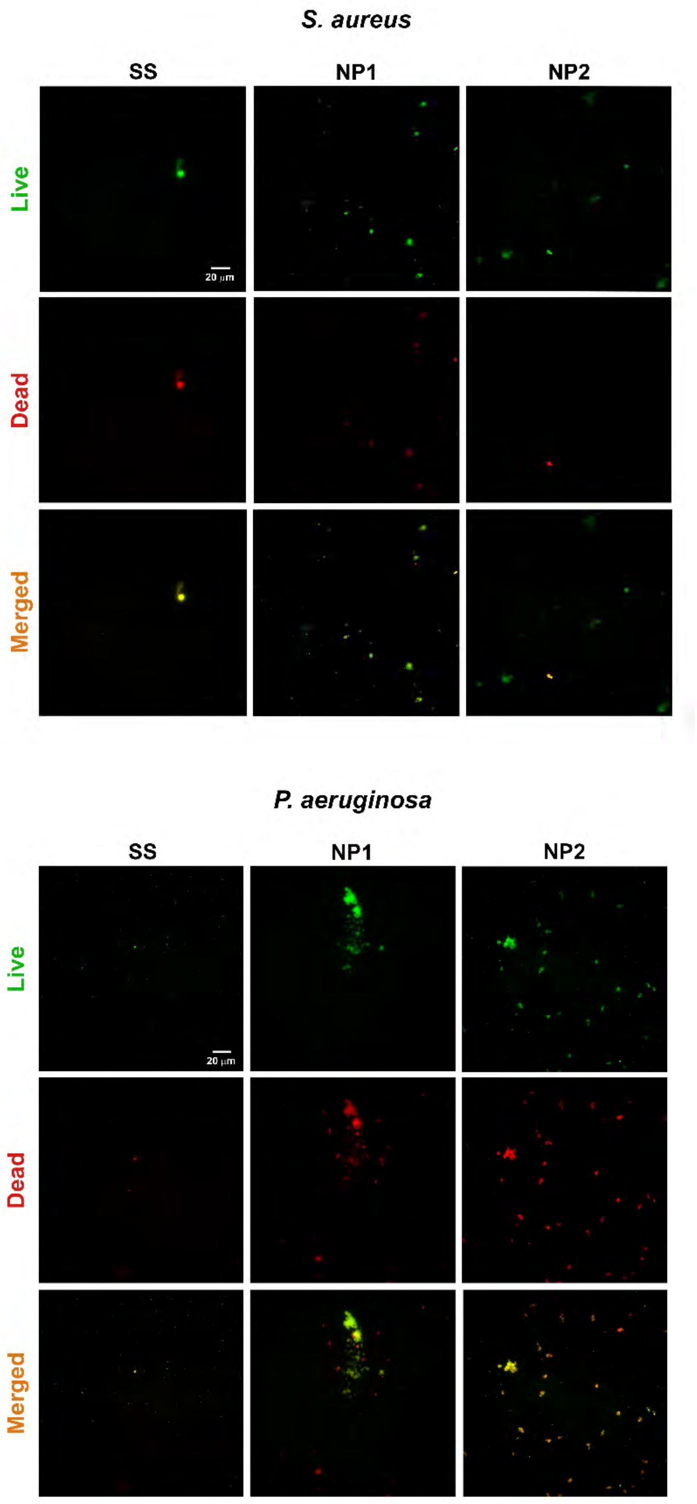


**Figure SI7.** Bacterial viability of *S. aureus* and *P. aeruginosa* on SS, NP1, and NP2. The live and dead bacterial cells were stained at 60 min using a live/dead assay, and representative images after cell staining were observed using a fluorescent microscope.
